# Comparison of microbiome and culture techniques for determination of gastrointestinal microbial communities in ceca of chickens

**DOI:** 10.1101/494781

**Authors:** Kim M. Wilson, Whitney N. Briggs, Audrey F. Duff, Kaylin M. Chasser, Xiaolun Sun, Lisa R. Bielke

## Abstract

The use of 16S next generation sequencing (NGS) technology to identify the relative abundance of microbial communities have become the standard when studying the intestinal microbiome. The increased use is due to the ability to identify a proportion of bacteria that cannot be observed with culture-based methods. However, culture-based techniques are acceptable to identify key bacterial groups, yet may grossly underestimate the microbial community in question. Since there is limited research comparing NGS results to colony forming units (CFU), the objective of this study was to compare total *Enterobacteriaceae* and lactic acid bacteria (LAB) recovery with culture techniques (CFU/g ceca) to total number of reads from operational taxonomic units (OTU) categorized as *Enterobacteriaceae* or LAB from Illumina MiSeq platform from matched chick cecal samples at three and 10 days of age. Both CFU recovery (1.09×10^9^ ± 2.42×10^8^; 1.37×10^8^ ± 5.57×10^7^) and reads (5460 ± 1164; 282 ± 163) belonging to *Enterobacteriaceae* decreased by 10 days of age (p < 0.001). Similarly, LAB reads decreased over time (21,128 ± 2262; 6220 ± 817, respectively p < 0.0001). However, LAB CFU recovery increased by 10 days (1.18×10^8^ ± 1.91×10^7^; 1.62×10^9^ ± 5.00×10^8^, respectively p < 0.01). At three days the Pearson’s correlation was -0.082 between CFU of culturable *Enterobacteriaceae* to reads and culturable LAB CFU to reads at 0.097, showing no correlation (p = 0.606, 0.551; respectively). By 10 days, no correlation of reads and CFU occurred with *Enterobacteriaceae* (r=-0.049; p-value = 0.769) while with LAB the correlation was 0.290 (p = 0.066) at 10 days. The CFU may be appropriate to identify a few families that change due to treatment or product. Without identifying viable cells to DNA recovered from NGS, there will always be the question whether the reads within the binned OTU in the intestinal tract is accurate.

## Introduction

Ascertaining bacterial levels in samples is among one of the most fundamental procedures in microbiology. Historically, measuring CFU on agar plates has been the most common method, though quantitative PCR and other gene-based methods, such as next-generation sequencing (NGS) have developed rapidly in the past 25 years [1–3]. The CFU enumeration method has disadvantages that include miscounting due to clumping of cells and inability to grow some types of bacteria in culture, commonly referred to as viable but non-culturable species. Often with these species, microbiologists are unable to decipher and replicate essential aspects of the gastrointestinal tract (GIT) environment within *in vitro* conditions. This limits the ability to grow some species of bacteria, especially those that grow within micro-environments of the GIT or those that are dependent on metabolic by-products and may require co-culture with other species [4].

Recent advances in accessibility and affordability of NGS and metagenomics techniques have increased its appeal to scientists that study microbial populations of the GIT because it has provided an ability to measure bacteria beyond only those that could be detected by culture methods and PCR. Metagenomic analysis requires reference-based identification and quantification by aligning the reads However, there are no current standardized approaches for these tools, as our understanding of the microbiome within the gut is at its infancy, especially as scientists try to understand what is or is not biologically relevant [5]. Thus, it is essential to understand variations of the microbiome in response to dietary amendments, additive insertion, litter management, etc. in relation to gut health [6–8]. Abundance taxonomy does not supply the complete picture required to understand how any treatment impacts the health of the GIT or the causation of dysbacteriosis, known to have detrimental effects on the host, because it does not include host responses, nor does it present insight into phenotypic changes within the microbiome [9–11]. However, there may be circumstances under which culture methodologies are sufficient, such as measuring changes in the levels of *Salmonella* within ceca. Though, culture-based techniques of digesta typically allow for differentiation at the genera level, full identification often requires multiple biochemical tests, and still often relies upon gene-based techniques to confirm identification. The objective of this study was to compare culture recovery of bacteria on MacConkey agar (MAC), for recovery of Gram-negative bacteria, and De Man, Rogosa and Sharpe agar (MRS) for recovery of lactic acid bacteria (LAB), to paired NGS microbiome detection of bacteria within the ceca of chickens. The goal of these comparisons was to determine differences and similarities of these methodologies in order to help scientists assess appropriate methods for their research goals while studying microbiota.

## Materials and Methods

### Embryo incubation and animal housing

Fertile broiler eggs were obtained from a local hatchery and placed in a single-stage incubator (Natureform Inc. Jacksonville, FL) until 18 embryonic days (18ED), then were randomly placed into one of 12 benchtop hatchers (Hova-Bator model 1602N, Savannah, GA, USA), which had been disinfected with 10% bleach prior to use. Up to 30 eggs were placed each hatcher to avoid overcrowding.

Immediately post-hatch, 128 chicks were placed into brooder battery cages that had wire floors covered with paper to encourage exposure of chicks to fecal material, and the paper was allowed to disintegrate naturally for the duration of the experiment. Age-appropriate ambient temperature was maintained with 24 hour lighting during the first seven days, followed by one hour of darkness through 10 days. Chicks has *ad libitum* access to a standard corn soy diet and water [12]. All activities were conducted under approved Institutional Animal Care and Use Committee protocols.

### Sample Collection

At three days post hatch, 80 chicks were removed for culture and microbiome analysis, with two chicks pooled to create a single sample (n=40). Since chicks were substantially larger with more digesta content, at 10 days, 45 chicks were sampled for both microbiome and culture analysis. Chicks were killed via cervical dislocation and all samples were aseptically collected post mortem. From each chick, a cecum was designated for culture-based recovery of bacteria, and the other for microbiome NGS.

### Culture Isolation and Enumeration

Cecal samples were placed into sterile bags and pulverized with a rubber mallet to expose contents. Samples were then weighed and 0.9% sterile saline added to make a five-fold dilution, after which serial 10-fold dilutions were made for plating on MAC (VWR, Suwanee, GA, USA) and MRS (Difco^TM^ Lactobacilli MRS AGAR VWR, Suwanee, GA, USA). All agar plates were incubated for up to 24 h at 37°C in an aerobic environment. Enumeration values from each selective medium for each sample were calculated to the original CFU/gram of ceca.

### DNA Isolation and Library Preparation

Ceca for DNA isolation were placed in 1.5mL tubes and flash-frozen in liquid nitrogen at the time of collection, then stored at −80 °C until further use. The Qiagen stool kit (Qiagen, Valencia, CA, USA) was utilized for DNA isolation following methods of Yu and Morrison [13]. Quantification of extracted DNA was completed with a Synergy HTX multi-mode plate reader (BioTek U.S., Winooski, VT, USA) and 5ng of DNA from each sample were randomly aliquoted onto a 96-well plate. All plates included known bacterial isolates as a positive control, plus a well with milliQ water as a negative control. MiSeq library preparation and 2 x 300 paired-end sequencing (Illumina, San Diego, CA, USA) was performed by the Ohio State University Molecular and Cellular Imaging Center (OSU MCIC). Primers amplified the V4-V5 region of the 16S rRNA gene (515F: GTGYCAGCMGCCGCGGTAA, 806R: GGACTACHVGGGTWTCTAAT; Ballou et al., 2016).

### 16S Sequence Data Analysis

After quality control from sequencing a total of 36,437,234 Illumina MiSeq reads were generated, and represented the prepared sequencing libraries for this study. The raw FASTQ files were de-multiplexed, primers and spacers removed and quality-filtered with QIIME 1.9.0 [15]. Quality filter variables included a minimum length of 200 base pairs, average quality score of 20, no barcode errors, reads were removed if there were two nucleotide mismatched, and chimeras were removed. Similar sequences were clustered together using the open-reference operational taxonomic unit (OTU) picking protocol, sequences were grouped into OTUs based on 97% sequence identity using the Ribosomal Database Project, with a minimum of 1,000 reads for each sample. Singleton and doubleton OTUs were removed due to potential sequencing errors or non-significant microbes. After initial quality processing, a total of 339 OTUs were identified. Represented sequences from each OTU were picked and assigned taxonomy and sequences with high identity (> 97%), and were clustered in the same OTU. The sequence coverage was normalized across all samples with a log_10_ transformation. Taxonomic assignments were generated using QIIME and R-studio (RStudio, Inc.). Taxonomic affiliations of sequences or reads classified under family *Enterobacteriaceae* or order Lactobacillales were counted which indicated the total number of reads within each binned OTU identified as either *Enterobacteriaceae* family or Lactobacillales order. These classifications were chosen because they primarily cover the types of bacteria recovered on MAC and MRS.

### Statistical Analysis

A Student’s t-test was performed to identify if there were changes in the number of reads and CFU/g ceca of *Enterobacteriaceae* and LAB over time, three days and 10 days (JMP 12.2.0, SAS Institute Inc). Observational jitter boxplots were constructed to visualize the distribution of cecal CFU/g of ceca for every sample from the MAC and MRS, as well as the number of reads belonging to OTUs categorized as *Enterobacteriaceae* or LAB on each day of collections (RStudio, Inc.). To evaluate the relationship between number of reads and CFU, a simple linear regression model was developed based on the equation:

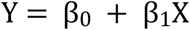

With β_0_ representing the intercept, and β_1_ representing the slope in relation to the corresponding to X or the CFU/g ceca count. The data were fitted to a straight line by the least squares method, which assessed the relationship in the model using the coefficient of determination, R^2^ and the relationship between the two variables using Pearson’s correlation coefficient ρ, and a p-value of < 0.05 were deemed significant (JMP 12.2.0, SAS Institute Inc).

## Results and Discussion

The use of culture-independent methods to characterize microbial communities has increased astronomically over the past few years, both due to major advances in NGS and ability to identify a large proportion of bacteria that are difficult or impossible to observe with culture based techniques [16]. Both culture dependent and culture independent methods have a role in microbiome studies; however, depending on what is being assessed one method may be better than the other or they should be used together. This research study compared culture and NGS methods to elucidate differences and similarities of the methods to allow for better use of these technologies in avian microbiome research.

Total reads from binned OTU *Enterobacteriaceae* and *Lactobacillales* at three days of age were significantly (p <0.0001) higher (1,063,533) compared to the total number of reads (266,582) present in the ceca at 10 days of age. The genus *Pseudomonas* can grow on MAC and is described as colorless [17]. *Pseudomonas* was not present with culture methods or microbiome analysis, thus the family *Enterobacteriaceae* alone was analyzed. At the neonatal stage of any mammalian or avian species, the diversity and abundance of microflora is generally low and as the animal ages this diversity and abundance increases. The increased intestinal diversity inadvertently decreases the relative abundance of predominantly aerobic or facultative anaerobic bacteria including the *Lactobacillales* order and *Enterobacteriaceae* family [14,18–20]. In this study, the *Enterobacteriaceae* populations significantly decreased by 10 days of age according to OTU counts (5460 ± 1164; 282 ± 163, p < 0.0001) and CFU recovery (1.09×10^9^ ± 2.42×10^8^; 1.37×10^8^ ± 5.57×10^7^, p < 0.001), respectively. Similarly, LAB OTU counts showed a decrease over time from three to 10 days of age (21,128 ± 2262 at three days of age; 6220 ± 817 at 10 days of age, p < 0.0001). However, LAB recovery significantly increased in CFU from three to 10 days of age (1.18×10^8^ ± 1.91×10^7^ at three days of age; 1.62×10^9^ ± 5.00×10^8^ at 10 days of age, p < 0.01). Figure 1 shows the differences seen in relative abundance from three to 10 days of age. The relationship between CFU and OTU did not correlate at three days of age (Fig 1A). At this time, the total recovered *Enterobacteriaceae* on MAC represented 64.8% of the total relative CFU/g and the remaining 35.2% belonged to LAB (p < 0.001); however, at the OTU level, *Enterobacteriaceae* represented 22.2% of the total relative reads and the remaining 77.8% belonged to the LAB (p < 0.01). By 10 days of age, the proportions were not significantly different in the culture and NGS methods with *Enterobacteriaceae* represented at 8.1% and 4.6% of the CFU/g ceca and OTU counts, respectively, while LABs accounted for 91.9% and 95.4% of the CFU and OTU counts, respectively (Fig 1B). The CFU/g ceca and total number of reads data were analyzed separately to provide the relationship of bacterial presence over time with the two types of techniques. However, because CFU and reads from binned OTUs do not provide the same units of measure, they could not be directly analyzed for statistical relevance. The differences between CFU and OTU at three days of age in Fig 1A may reflect the rapid turnover of bacteria that could be dormant, non-viable or non-culturable.

**Fig. 1.**
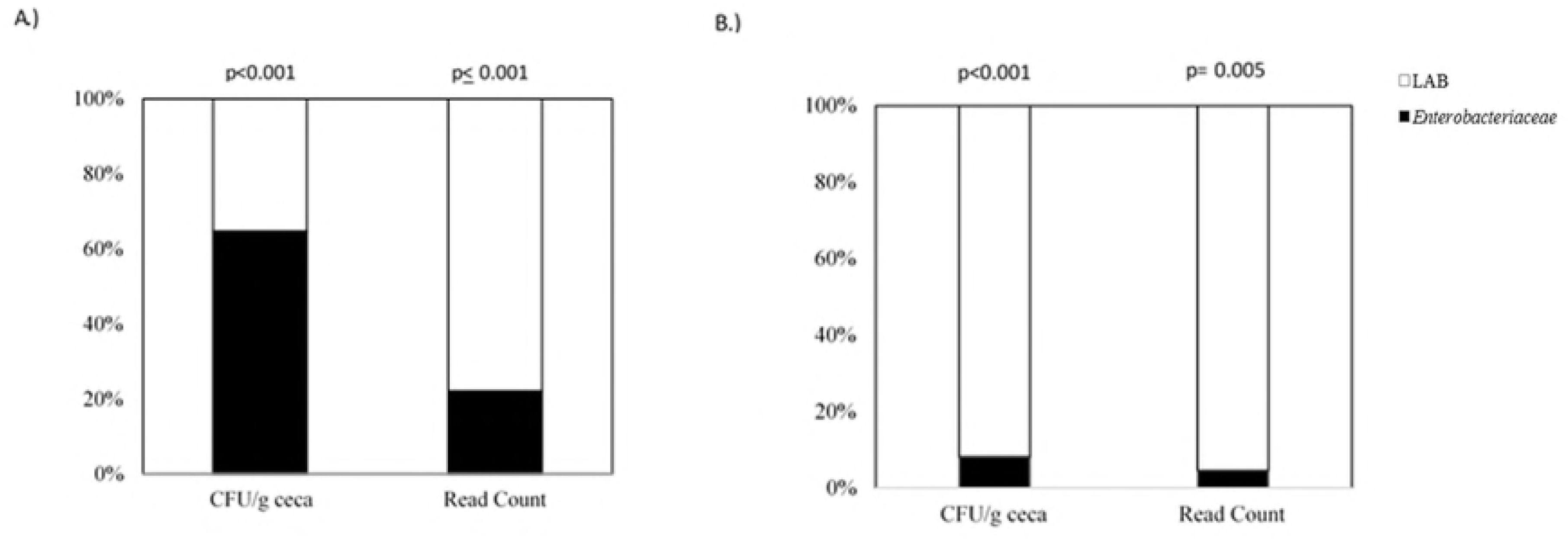
Proportion of *Enterobacteriaceae* and LAB CFU/g ceca or Reads from respective *Enterobacteriaceae* and LAB OTU. (A) three days (n=40); (B) 10 days (n=45) post-hatch. The p-value indicates comparison of *Enterobacteriaceae* and LAB within methodology (CFU/gram ceca or total reads from binned OTU) and time of collection, three days and 10 days.

The retention time of feed in the small intestine has been determined to be approximately three hours, with the ileum being the bottleneck as there are a reduced amount of digestible components in this space that can slow the flow rate [21]. Between four and 10 days of age, broilers have been observed to experience a rapid increase in feed intake with a parallel decrease in passage rate, which may be partially due to quicker absorption of amino-acid derived nitrogen and fatty acids [22]. This change in digesta flow may account for nutrient availability for bacteria at three days of age. For example, *Lactobacillus* are a complex organism that require certain amino acids, vitamins and sugars to thrive [23].

Results in this experiment show that reads from OTUs do not provide the same answers at CFU recovery for particular times of collection. Rubinelli et al (2017) found the inclusion of acidifiers such as sodium bisulfate (SBS) were associated with decreased recovery of *Salmonella* from *in vitro* spiked chicken cecal contents at 48 hours post incubation; however, the relative abundance showed no difference in microbial composition after 24 hours of incubation and no difference of the abundance of Gammaproteobacteria class or *Enterobacteriaceae* family, in which *Salmonella* belongs [24]. For this study, the viable bacteria recovered on MAC ranged from 3.50×10^7^ to 8.00×10^9^ CFU/g ceca at three days of age (Fig 2A).

**Fig 2.**
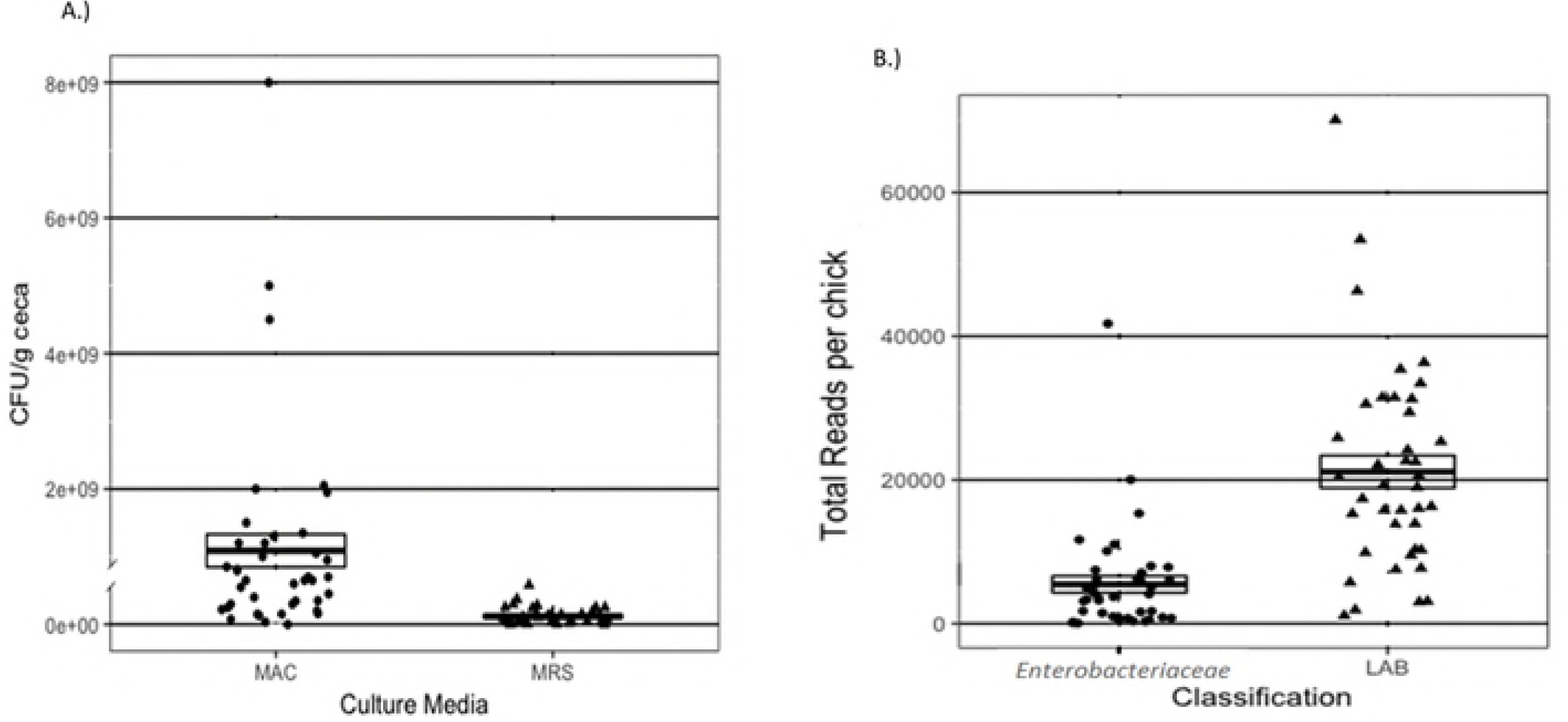
Jitter boxplot representing total CFU/g ceca or Reads from Enterobacteriaceae or LAB genera at three days. (A) CFU/g ceca from MacConkeys agar (MAC), Man Rogosa and Sharpe agar (MRS), (B) total number of reads from the *Enterobacteriaceae* family or lactic acid producing bacteria (LAB) order Lactobacillales OTUs from cecal samples (n=40) at three days of age. Each dot represents a single sample. The solid black line represents the mean. The box outline represents the ± standard deviation

Conversely, *Enterobacteriaceae* OTU number of reads had a distribution of 63 to 41,741 total reads at 3 days of age (Fig 2B). By 10 days of age, *Enterobacteriaceae* MAC recovery range was 2.50×10^5^ to 2.00×10^9^ CFU/g ceca (Fig 3A) and number of reads had a range of 0 to 6,327 OTU counts (Fig 3B). The LAB recovery distribution at 3 days of age was 0.00 to 5.80×10^8^ CFU/g on MRS (Fig 2A) whereas the number of reads from LAB OTU ranged from 1,105 - 70,087 (Fig 2B). By 10 days of age, the distribution of LAB recovered on MRS was 1.30×10^7^ to 1.5×10^10^ CFU/g ceca (Figure 3A) and the OTU counts had a range of 207 to 21,105 (Fig 3B). The results showed no correlation between total reads and CFU counts for *Enterobacteriaceae* (ρ =-0.082; p-value = 0.606; Fig 4A) or LAB (ρ = 0.097; p-value = 0.551; Fig 4B) and no correlation at 3 days of age (ρ = -0.049; p-value = 0.769; Fig 5A) or 10 days or age (ρ = 0.290; p-value = 0.066; Fig 5B). Currently, there are no standard methods for comparison of CFU and OTU data. These figures (Fig 1-5) illustrate the different patterns within the same day and type of bacteria recovered.

**Fig. 3.**
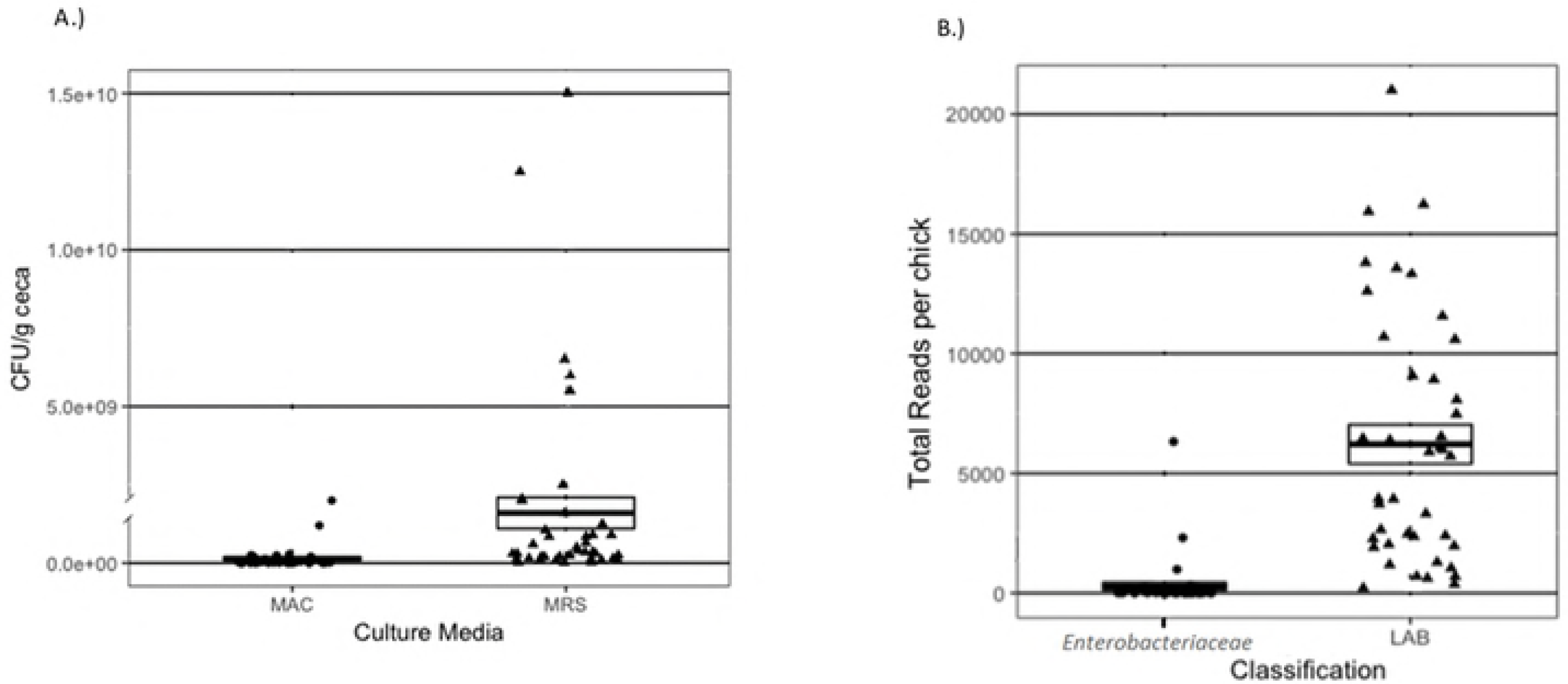
Jitter boxplot representing total CFU/g ceca or Reads from Enterobacteriaceae or LAB genera at ten days. (A) CFU/g ceca from MacConkeys agar (MAC), Man Rogosa and Sharpe agar (MRS), (B) total number of reads from the *Enterobacteriaceae* family or lactic acid producing bacteria (LAB) order Lactobacillales OTUs from cecal samples (n=45) at three days of age. Each dot represents a single sample. The solid black line represents the mean. The box outline represents the ± standard deviation

**Fig. 4.**
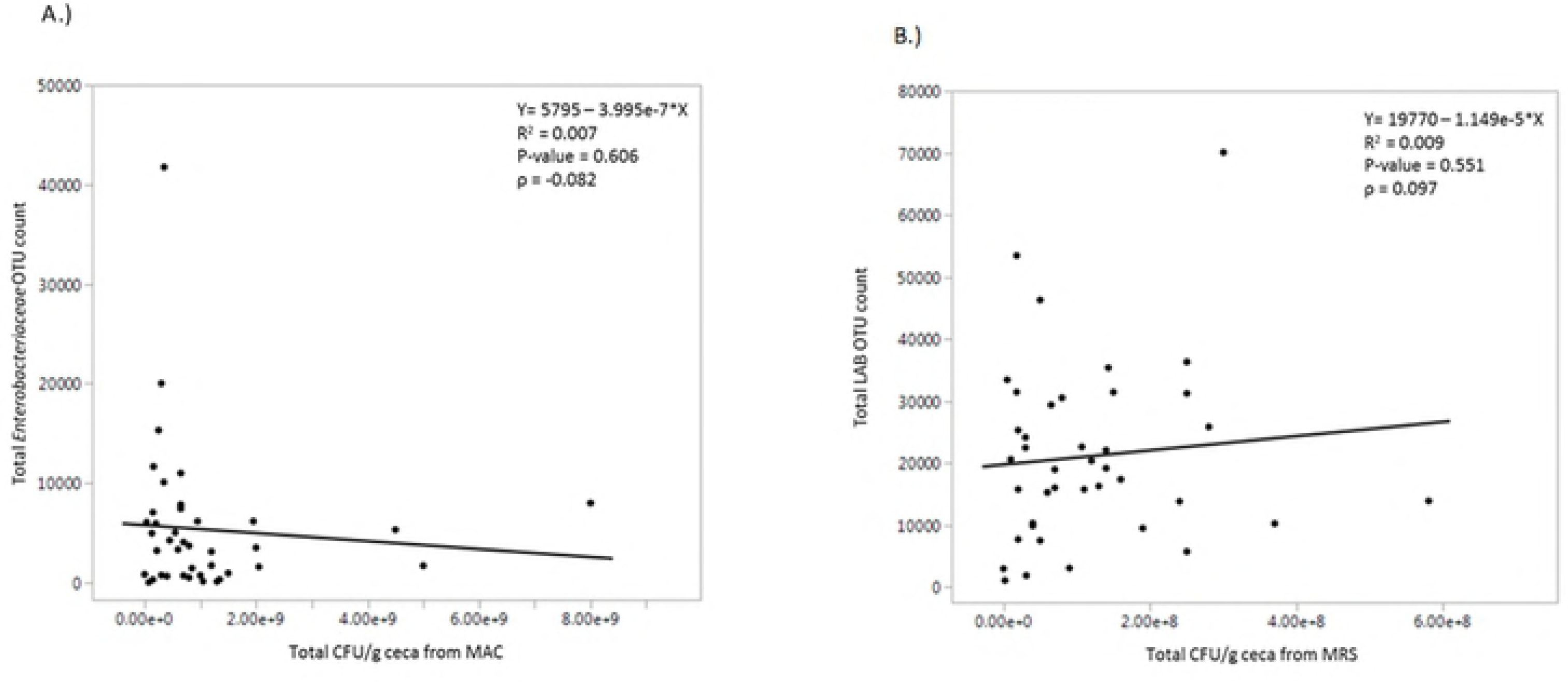
Dot plot depicting the relationship of total number of reads and CFU/g ceca of A) *Enterobacteriaceae* and B) LAB from three day-old chicks (n=40). The black line represents the fitted regression linear regression line with its corresponding equation and correlation of determination (R^2^), Pearson’s correlation coefficient between number of reads and CFU (ρ) and the p-value

**Figure 5.**
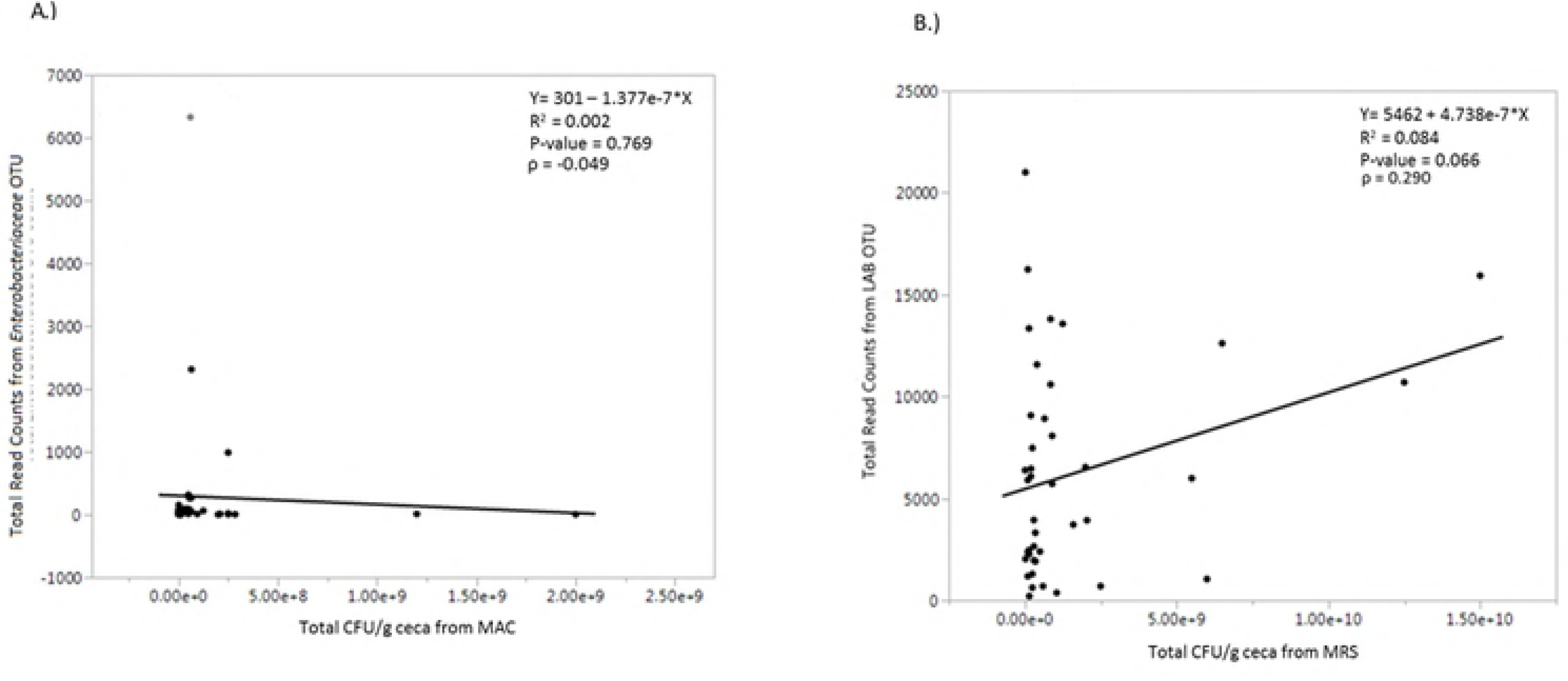
Dot plot depicting the relationship of total number of reads and CFU/g ceca of A) *Enterobacteriaceae* and B) LAB from ceca of 10 day old chicks (n=45). The black line represents the fitted regression linear regression line with its corresponding equation and correlation of determination (R^2^), Pearson’s correlation coefficient between number of reads and CFU (ρ) and the p-value

Quantifying microbiota by 16S rRNA have not been able to determine associations with microorganisms in an active or potentially active state because the results include active as well as dormant, dead and quiescent bacteria which are present in all microbial samples [25]. This lack of ability to detect DNA from only active cells beg the question whether the number of reads within the binned OTUs in any biological system are accurate representations of active microbial communities. In this study, the total relative abundance of *Enterobacteriaceae* and LABs were different between three and 10 days of age (three days of age: 4.24%, 39.37%; 10 days of age: 21.74%, 1.91%, respectively; data not shown). In general, the alpha diversity (i.e. number of species present within a sample) is increased as the animal ages and it is unlikely that culture based recovery will be able to reflect this change since a wide variety of media would be required and viable but non-culturable species would not be detected [14,20,26,27]. Many *Lactobacillales* are facultative anaerobes but some species, such as *Lactobacillus fermennti*, are slow growing and other genera, such as *Leuconostoc* and *Pediococcus*, may not grow as fastidiously as *Lactobacillus* on MRS agar [28].

The bacterial groups recovered from the *Enterobacteriaceae* family included the genus *Citrobacter, Klebsiella* (i.e. *Citrobacter* 2), *Proteus* and *Trabulsiella*. The total OTUs present in the LAB or *Lactobacillales* order included the genus *Enterococcus, Lactobacillus, Pediococcus, Streptococcus* and *Leuconostoc*. A common marker that researchers examine in improved gut health is an increase LAB. Many bacteria produce lactic acid as an end-product, a LAB primary metabolic end product is lactic acid by glucose metabolism. The LAB are functionally comprised of catalase negative and Gram-positive bacteria and the group typically falls under the order *Lactobacillales* [29]. The LABs include the genus *Lactobacillus, Enterococcus, Lactococcus, Leuconstoc, Pediococcus* and *Streptococcus*. Choosing known families or orders of bacteria such as *Enterobacteriaceae* and *Lactobacillales* to monitor with culture based techniques may be sufficient to answer questions related to efficacy of a treatment since the counts are of live culturable bacteria and not DNA.

The term CFU is based on the idea that microbial colonies counted on agar began as a single cell or aggregates of several cells [30]. In DNA-based sequencing techniques, the number of taxonomic units that have been identified in a given sample are deemed as OTU. This unit is defined based on sequence similarity [31]. For this similarity, 97% of the DNA sequence of the bacterial 16S rRNA gene is commonly used to define a taxonomic genus and 98-99% for species [31]. This level of sequence resemblance is believed to compare the common species but this can be based on morphology, physiology and other characteristics of organisms, so the terms species and OTU may not be fully comparable [31].

Some researchers support their 16S rRNA sequencing with previous CFU findings, particularly for a single bacterial genus [32,33]. However, a consequence of the immense information that microbiome-based analysis provides is there may be more difficulty to find statistically different data compared to CFU counts [34]. Approximately 20% of bacteria in the gastrointestinal tract (GIT) of a chicken are culturable so CFU counts alone will provide an underestimation of the total microbiota present [35]. Analysis of the microbial composition is not necessarily limited by technical abilities, but rather inconsistencies throughout the literature and experiments, including bias generated by PCR, as well as sequence coverage and length, which greatly alter the taxonomic information [25,36–39].

The increased high throughput abilities of NGS has provided researchers with invaluable information regarding the presence or absence of total bacteria in a microbial community. However, this technology does have limitations. This results from this study showed that culturing and NGS techniques demonstrated little to no correlation on the presence or absence of two groups of bacteria that were tested. While CFU/g ceca and OTU provide different results, each technique may be appropriate for different situations. For example, CFU would be acceptable when the objective is to identify culturable bacteria or recovery of a single or few species of known bacteria. Conversely, NGS is beneficial to identify overall community changes in situations where global DNA content may provide a more complete answer. Each methodology of quantification may yield different results so care should be taken to select the correct microbiological techniques to test a hypothesis in order to avoid drawing conclusions that are not represented by the type of data analyzed.

## Acknowledgements

The authors would like to thank the farm personnel at the OARDC Poultry Research Farm including Keith Patterson for providing the eggs. Kendal Searer is greatly acknowledged for her technical assistance as well as Saranga Wijerante, MS for his bioinformatics guidance. Lastly, the authors would like to acknowledge and thank the OARDC Research Enhancement Competitive Grants Program (SEEDS) #2016035 for providing the funding of this research

## Author Contributions

Conceived and designed the experiments: KMW, LRB. Participated in sample collections: KMW, WRB, AFD, KMC. Analyzed the data: KMW, XS. Contributed to reagents/materials/analysis tools: KMW, WRB, XS. Wrote the paper: KMW, LRB. Critical review of manuscript: LRB and XS.

## References

1. Monod J. The Growth of Bacterial Cultures. Annual Review of Microbiology. 1949;3: 371–394. doi:10.1146/annurev.mi.03.100149.002103

2. van Dijk EL, Auger H, Jaszczyszyn Y, Thermes C. Ten years of next-generation sequencing technology. Trends in Genetics. 2014;30: 418–426. doi:10.1016/j.tig.2014.07.001

3. Hagemann IS. Chapter 1 - Overview of Technical Aspects and Chemistries of Next-Generation Sequencing. In: Kulkarni S, Pfeifer J, editors. Clinical Genomics. Boston: Academic Press; 2015. pp. 3–19. doi:10.1016/B978-0-12-404748-8.00001-0

4. Stewart EJ. Growing Unculturable Bacteria. J Bacteriol. 2012;194: 4151–4160. doi:10.1128/JB.00345-12

5. He Y, Caporaso JG, Jiang X-T, Sheng H-F, Huse SM, Rideout JR, et al. Stability of operational taxonomic units: an important but neglected property for analyzing microbial diversity. Microbiom. 2015;3: 20. doi:10.1186/s40168-015-0081-x

6. Park SH, Perrotta A, Hanning I, Diaz-Sanchez S, Pendleton S, Alm E, et al. Pasture flock chicken cecal microbiome responses to prebiotics and plum fiber feed amendments. Poult Sci. 2017;96: 1820–1830. doi:10.3382/ps/pew441

7. Roto SM, Kwon YM, Ricke SC. Applications of *In Ovo* Technique for the Optimal Development of the Gastrointestinal Tract and the Potential Influence on the Establishment of Its Microbiome in Poultry. Front Vet Sci. 2016;3. doi:10.3389/fvets.2016.00063

8. Stanley D, Denman SE, Hughes RJ, Geier MS, Crowley TM, Chen H, et al. Intestinal microbiota associated with differential feed conversion efficiency in chickens. Appl Microbiol Biotechnol. 2012;96: 1361–1369. doi:10.1007/s00253-011-3847-5

9. Teirlynck E, Gussem MDE, Dewulf J, Haesebrouck F, Ducatelle R, Immerseel FV. Morphometric evaluation of “dysbacteriosis” in broilers. Avian Pathology. 2011;40: 139–144. doi:10.1080/03079457.2010.543414

10. Belkaid Y, Hand T. Role of the Microbiota in Immunity and inflammation. Cell. 2014;157: 121–141. doi:10.1016/j.cell.2014.03.011

11. Lan Lin, Jianqiong Zhang. Role of intestinal microbiota and metabolites on gut homeostasis and human diseases. BMC Immunology. 2017;18: 1–25. doi:10.1186/s12865-016-0187-3

12. Nutrient Requirements of Poultry [Internet]. Washington, D.C.: National Academies Press; 1994. doi:10.17226/2114

13. Yu Z, Morrison M. Improved extraction of PCR-quality community DNA from digesta and fecal samples. BioTechniques. 2004;36: 808–812.

14. Ballou AL, Ali RA, Mendoza MA, Ellis JC, Hassan HM, Croom WJ, et al. Development of the Chick Microbiome: How Early Exposure Influences Future Microbial Diversity. Front Vet Sci. 2016;3: 1–12. doi:10.3389/fvets.2016.00002

15. Caporaso JG, Lauber CL, Walters WA, Berg-Lyons D, Huntley J, Fierer N, et al. Ultra-high-throughput microbial community analysis on the Illumina HiSeq and MiSeq platforms. ISME J. 2012;6: 1621–1624. doi:10.1038/ismej.2012.8

16. Zapka C, Leff J, Henley J, Tittl J, De Nardo E, Butler M, et al. Comparison of Standard Culture-Based Method to Culture-Independent Method for Evaluation of Hygiene Effects on the Hand Microbiome. mBio. 2017;8. doi:10.1128/mBio.00093-17

17. Abbas M, Emonet S, Köhler T, Renzi G, van Delden C, Schrenzel J, et al. Ecthyma Gangrenosum: *Escherichia coli* or *Pseudomonas aeruginosa*? Front Microbiol. 2017;8. doi:10.3389/fmicb.2017.00953

18. Palmer C, Bik EM, DiGiulio DB, Relman DA, Brown PO. Development of the Human Infant Intestinal Microbiota. PLOS Biology. 2007;5: e177. doi:10.1371/journal.pbio.0050177

19. Houghteling PD, Walker WA. Why is initial bacterial colonization of the intestine important to the infant’s and child’s health? J Pediatr Gastroenterol Nutr. 2015;60: 294–307. doi:10.1097/MPG.0000000000000597

20. Awad WA, Mann E, Dzieciol M, Hess C, Schmitz-Esser S, Wagner M, et al. Age-Related Differences in the Luminal and Mucosa-Associated Gut Microbiome of Broiler Chickens and Shifts Associated with *Campylobacter jejuni* Infection. Front Cell Infect Microbiol. 2016;6: 1–17. doi:10.3389/fcimb.2016.00154

21. Liu JD, Secrest SA, Fowler J. Computed tomographic precision rate-of-passage assay without a fasting period in broilers: More precise foundation for targeting the releasing time of encapsulated products. Livestock Science. 2017;200: 60–63. doi:10.1016/j.livsci.2017.04.006

22. Noy Y, Sklan D. Digestion and absorption in the young chick. Poultry Science. 1995; 366–373.

23. Apajalahti J, Vienola K. Interaction between chicken intestinal microbiota and protein digestion. Animal Feed Science and Technology. 2016;221, Part B: 323–330. doi:10.1016/j.anifeedsci.2016.05.004

24. Rubinelli PM, Kim SA, Park SH, Roto SM, Ricke SC. Sodium bisulfate and a sodium bisulfate/tannin mixture decreases pH when added to an in vitro incubated poultry cecal or fecal contents while reducing *Salmonella* Typhimurium marker strain survival and altering the microbiome. Journal of Environmental Science and Health, Part B. 2017;52: 607–615. doi:10.1080/03601234.2017.1316159

25. Rojo D, Méndez-García C, Raczkowska BA, Bargiela R, Moya A, Ferrer M, et al. Exploring the human microbiome from multiple perspectives: factors altering its composition and function. FEMS Microbiol Rev. 2017;41: 453–478. doi:10.1093/femsre/fuw046

26. Sergeant MJ, Constantinidou C, Cogan TA, Bedford MR, Penn CW, Pallen MJ. Extensive Microbial and Functional Diversity within the Chicken Cecal Microbiome. PLOS ONE. 2014;9: e91941. doi:10.1371/journal.pone.0091941

27. Oakley BB, Kogut MH. Spatial and Temporal Changes in the Broiler Chicken Cecal and Fecal Microbiomes and Correlations of Bacterial Taxa with Cytokine Gene Expression. Front Vet Sci. 2016;3: 1–12. doi:10.3389/fvets.2016.00011

28. Schillinger U, Lücke F-K. Identification of lactobacilli from meat and meat products. Food Microbiology. 1987;4: 199–208. doi:10.1016/0740-0020(87)90002-5

29. Salvetti E, Torriani S, Felis GE. The Genus Lactobacillus: A Taxonomic Update. Pro Antimicro Prot. 2012;4: 217–226. doi:10.1007/s12602-012-9117-8

30. Ben-Jacob E, Schochet O, Tenenbaum A, Cohen I, Czirók A, Vicsek T. Generic modelling of cooperative growth patterns in bacterial colonies. Letters to Nature. 1994;368: 46–49. doi:10.1038/368046a0

31. Rosselló-Mora R, Amann R. The species concept for prokaryotes. FEMS Microbiol Rev. 2001;25: 39–67. doi:10.1111/j.1574-6976.2001.tb00571.x

32. Schlaberg R, Chiu CY, Miller S, Procop GW, Weinstock G. Validation of Metagenomic Next-Generation Sequencing Tests for Universal Pathogen Detection. Archives of Pathology & Laboratory Medicine. 2017;141: 776–786. doi:10.5858/arpa.2016-0539-RA

33. Sun X, Winglee K, Gharaibeh RZ, Gauthier J, He Z, Tripathi P, et al. Microbiota-derived Metabolic Factors Reduce Campylobacteriosis in Mice. Gastroentero. 2018; doi:10.1053/j.gastro.2018.01.042

34. Olsen R, Kudirkiene E, Thøfner I, Pors S, Karlskov-Mortensen P, Li L, et al. Impact of egg disinfection of hatching eggs on the eggshell microbiome and bacterial load. Poult Sci. 2017;96: 3901–3911. doi:10.3382/ps/pex182

35. Gaskins HR, Collier CT, Anderson DB. Antibiotics as growth promotants: mode of action. Animal Biotechnology. 2002;13: 29–42.

36. Youssef N, Sheik CS, Krumholz LR, Najar FZ, Roe BA, Elshahed MS. Comparison of Species Richness Estimates Obtained Using Nearly Complete Fragments and Simulated Pyrosequencing-Generated Fragments in 16S rRNA Gene-Based Environmental Surveys. Appl Environ Microbiol. 2009;75: 5227–5236. doi:10.1128/AEM.00592-09

37. Stanley D, Hughes RJ, Moore RJ. Microbiota of the chicken gastrointestinal tract: influence on health, productivity and disease. Appl Microbiol Biotechnol. 2014;98: 4301–4310. doi:10.1007/s00253-014-5646-2

38. Keller A, Horn H, Förster F, Schultz J. Computational integration of genomic traits into 16S rDNA microbiota sequencing studies. Gene. 2014;549: 186–191. doi:10.1016/j.gene.2014.07.066

39. Jovel J, Patterson J, Wang W, Hotte N, O’Keefe S, Mitchel T, et al. Characterization of the Gut Microbiome Using 16S or Shotgun Metagenomics. Front Microbiol. 2016;7. doi:10.3389/fmicb.2016.00459

